# DrugDiff - small molecule diffusion model with flexible guidance towards molecular properties

**DOI:** 10.1101/2024.07.17.603873

**Authors:** Marie Oestreich, Erinc Merdivan, Michael Lee, Joachim L. Schultze, Marie Piraud, Matthias Becker

## Abstract

With the cost/yield-ratio of drug development becoming increasingly unfavourable, recent work has explored machine learning to accelerate early stages of the development process. Given the current success of deep generative models across domains, we here investigated their application to the property-based proposal of new small molecules for drug development. Specifically, we trained a latent diffusion model — *DrugDiff* — paired with predictor guidance to generate novel compounds with a variety of desired molecular properties. The architecture was designed to be highly flexible and easily adaptable to future scenarios. Our experiments showed successful generation of unique, diverse and novel small molecules with targeted properties. The code is available at https://github.com/MarieOestreich/DrugDiff.

## Main

The drug development process has become increasingly unsustainable over the last years, with the ratio of product yield to development costs becoming more and more unfavourable [1], [2], [3], [4]. In an attempt to revert this trend, we can observe the development of generative machine learning models in recent years [5], [6], [7], [8], [9], [10], [11], [12], [13]. Particular advancements have been made recently with diffusion models for AI-based protein design [7], [8], [11]. However, the drug space, i.e. the parts of the chemical space harvested for drug development, not only comprises proteins. The three dominant chemical subspaces are small molecules, proteins and nucleic acids such as RNA (Fig.1A). With small molecules making up the majority of drugs [14], we developed *DrugDiff*, a diffusion model for the generation of small molecules (Fig. 1B), to help navigate the vast space of potentially drug-like small molecules. Several aspects suggested diffusion models as suitable candidates for this task: **i)** their recent successes not only in the image but also the protein domain demonstrate their potential to model highly complex data; **ii)** when designing novel small molecule drugs, it is essential to offer guidance towards desired molecular properties during the generation process. To maximise utility and applicability in diverse scenarios, the set of properties to guide for must be highly flexible. With predictor-based guidance, diffusion models offer guided generation without explicit conditional training, introducing the required flexibility; **iii)** unlike other generative model architectures, diffusion models do multi-step rather than one-shot generation. This allows gradual guidance towards desired properties with the option to correct missteps along the way and **iv)** they can be combined with variational autoencoders (VAEs) to form a latent diffusion model. Latent diffusion models are not trained on the original data space, but instead on a latent representation stemming from a pre-trained VAE. Latent diffusion models are particularly attractive in the context of small molecules, to address the often-raised topic of molecule representation, because unlike proteins and nucleic acids, small molecules are not chains of pre-existing building blocks like amino acids or nucleobases that are connected in a clearly defined manner. Instead, there are complex chemical rules that dictate, for instance, each atom’s number of eligible bonds, possible charges or bond angles and these rules are difficult to represent. While several molecular representations exist [15], many of them are either discrete, not unique, or very sparse, whereas a continuous numeric representation is the preferred input for many deep learning models. Latent diffusion models outsource the task of learning a mapping from the molecular representation to a continuous latent space to a VAE, while the training of the diffusion model itself is focussed on modelling its latent distribution. Precisely, our proposed model *DrugDiff* comprises three parts (Fig. 1C): 1) A VAE trained on SELFIES representations of small molecules [16] from the public ZINC250K dataset, which contains approximately 250,000 small, drug-like and commercially available molecules from the ZINC database [17], [18]; 2) a latent diffusion model trained on the VAE’s latent space and 3) a series of molecular property predictors trained independently on one-hot encodings of the ZINC250K molecules. The diffusion model learns to generate latent representations of small molecules by starting from pure Gaussian noise and predicting — in a series of diffusion steps — small amounts of noise to remove and gradually de-noise the latent representation. To guide this diffusion process towards desired molecular properties, the latents are decoded at every step, the decoded molecules are then passed to the pre-trained property predictors and the computed loss between desired and actual property is back-propagated directly onto the latent space. The latent space is then manipulated using the computed gradients before entering the next denoising step. We chose this method to guide towards molecular properties because we decided to explicitly avoid conditional training of the diffusion model. The consequence of conditional training would be that whenever new properties are to be added in future applications, the diffusion model would have to be retrained. Not only would that make the model inflexible in its direct application to future use cases, but it would also render the model very unsustainable. We addressed these issues by using predictor guidance. To illustrate the successful generation of small molecules with defined molecular features with *DrugDiff*, we selected a variety of molecular properties that are relevant for the mechanism of action of drug-like small molecules (Fig. 1D).

**Figure 1:**
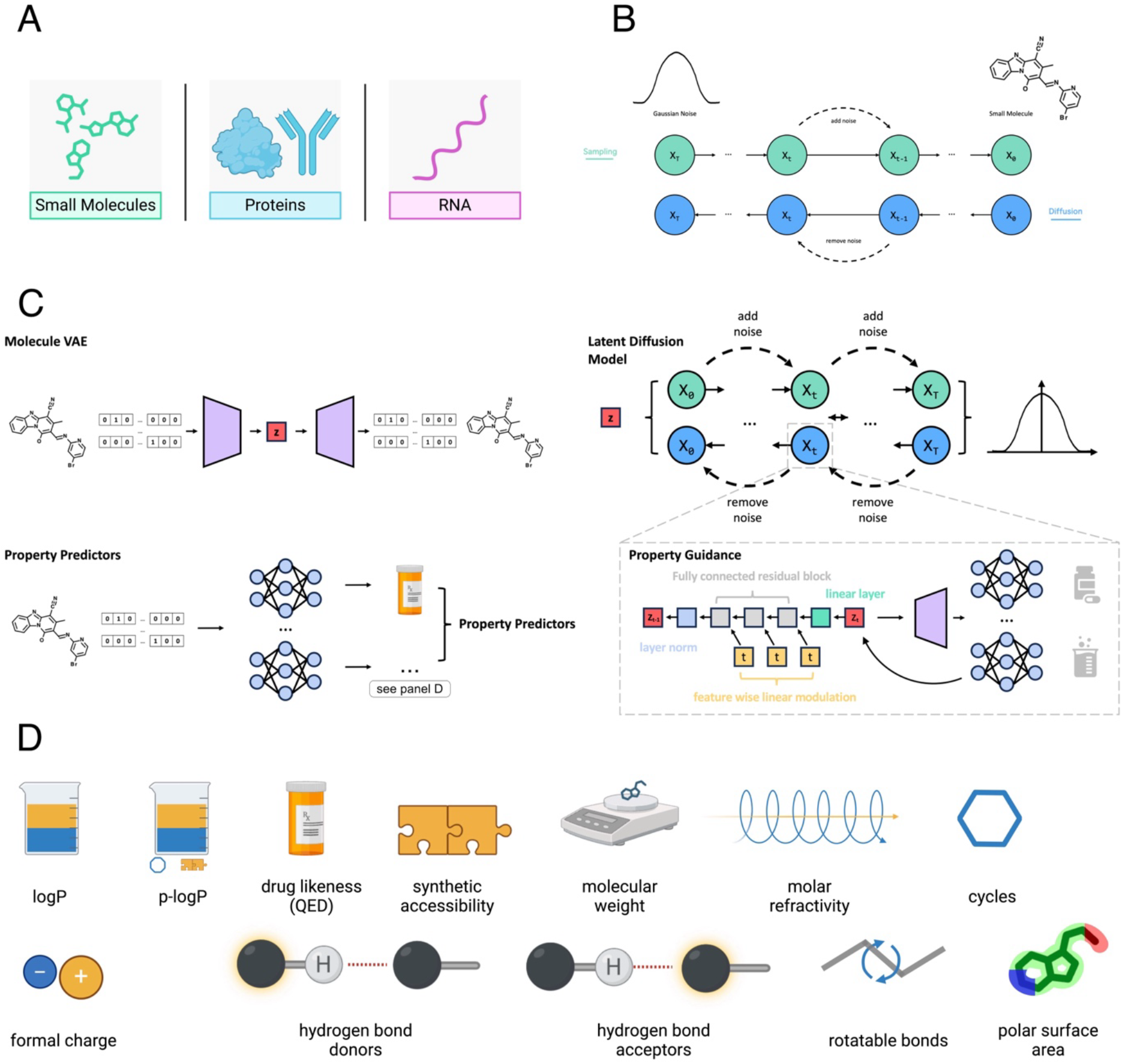
DrugDiff Overview. A) dominant chemical subspaces utilised for drug discovery; B) schematic of DrugDiff, which uses a diffusion model to generate molecules from Gaussian noise. Both the forward and the backward diffusion process are illustrated; C) the detailed architecture of DrugDiff, which comprises a VAE (top left) that autoencodes one-hot-encoded SELFIES and whose latent space (z) serves as input to the latent diffusion model (top right). The bottom left illustrates molecular property predictors that are trained on one-hot encoded SELFIES and then used for guidance during the diffusion steps. The guidance together with the detailed architecture of the latent diffusion model is illustrated in the inset (bottom right); D) The molecular properties used for guidance.

## Results

### Unguided Generation of Small Molecules

Before steering the generation process towards specific molecular properties, we first validated that the model had correctly learned from its training data distribution. To this end, we generated 10,000 molecules with *DrugDiff* without any guidance and additionally sampled the same number of molecules from the VAE’s latent space, the distribution that our diffusion model was trained to learn. Figure 2A shows a random subset of the molecules generated with *DrugDiff*. The molecules exhibit diverse structures, featuring various elements, ring sizes, bond types and molecular sizes. Figure 2B quantifies *DrugDiff*’s ability to produce unique, novel and chemically diverse molecules. The scores are generally high and comparable to that of the VAE. The internal diversity score measures how chemically diverse a set of molecules is on a scale from 0 (not diverse) to 1 (very diverse), with a high score of 0.91 indicating *DrugDiff* did not only learn subspaces of the latent space but covered its full chemical information landscape. The high novelty and uniqueness scores show that *DrugDiff* was able to generalise to the underlying data distribution and neither suffers from mode collapse nor overfitting. We additionally evaluated the VAE and *DrugDiff* using the GuacaMol benchmarking dataset and associated Distribution-Learning Benchmarks [19]. To this end, we re-trained the VAE on the benchmarking set and subsequently re-trained *DrugDiff* on the new VAE latent space. The benchmark evaluation (Suppl. Table 1) shows that the results for VAE and *DrugDiff* are very similar, further underlining that *DrugDiff* is indeed capable of fully learning the VAE’s latent space. We subsequently investigated in more detail whether *DrugDiff* could cover the full range of various molecular properties as they occurred in the training data. Figure 2C shows the distribution of 15 properties in the molecules generated with the VAE compared to those generated by *DrugDiff*. Those properties are: The topological polar surface area (a measurement for passive transport through membranes [20]), synthetic accessibility (a score to estimate how easily a molecule can be synthesised [21]), the number of rotatable bonds (single bonds that are not part of a ring and that are attached to an atom that is neither hydrogen nor terminal [22]), quantitative estimation of drug-likeness [23], molar refractivity (a measurement for how polarisable a molecule is [24]), molecular weight, the number of atoms and heavy atoms in particular, the logP (indicates a molecule’s lipophilicity [24]) as well as penalised logP (logP penalised for poor synthetic accessibility and large cycles [25]), the number of very small or large cycles (≤ 4 or ≥ 7 atoms), the number of hydrogen-bond donors and -acceptors, the number of cycles of any size, and formal charge. For all these properties, the property distributions of the molecules generated by *DrugDiff* and those randomly sampled from the latent space are highly overlapping.

**Figure 2:**
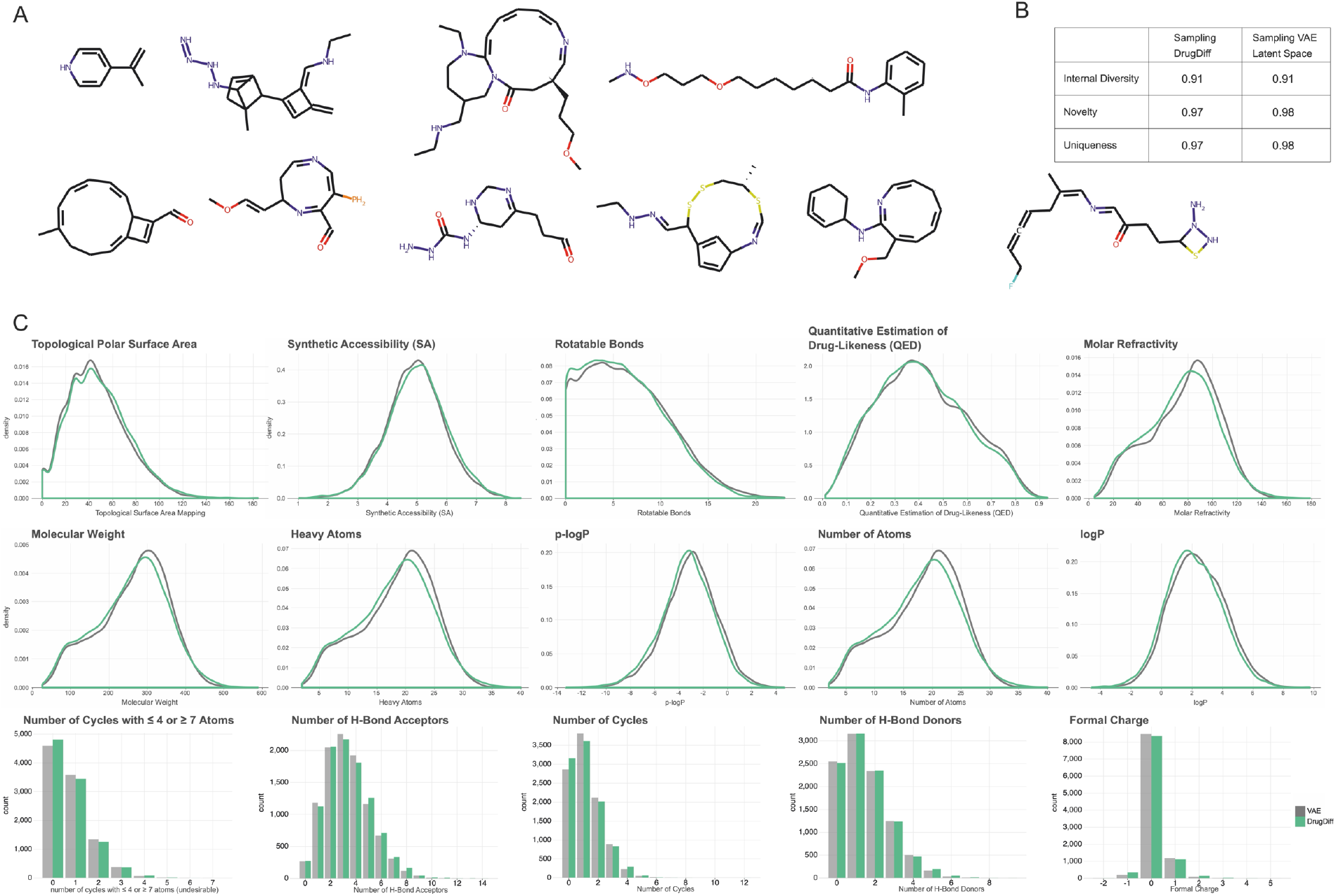
Unguided Generation. A) Randomly chosen subset of molecules generated by DrugDiff without guidance; B) Internal diversity, novelty and uniqueness of molecules generated with DrugDiff vs. directly sampled from the VAE latent space. All metrics are on a scale from 0 to 1 with 1 being the optimum; C) Distribution of different molecular properties in 10,000 molecules generated with DrugDiff (green) vs. directly sampled from the VAE latent space (grey).

In summary, these results illustrate that *DrugDiff* learned to cover the full distribution of the training data, and without guidance towards target properties it is capable of generating novel, unique and chemically diverse small molecules, demonstrating to good generalisation to the data space.

### Single-Property Guidance

For a model to truly facilitate drug development by proposing novel small molecules, it *must* be able to generate such molecules under provided property constraints. While unconditional generation with subsequent filtering for molecular properties is theoretically an option, the lack in targeted generation typically results in the loss of most generated molecules after filtering. Such an approach would require a much larger number of molecules to be generated to account for the filter loss and molecules with property values residing at the tails of their distributions will be underrepresented. Actively guiding the generative process towards a desired property is therefore important to maximise the yield of molecules in the acceptable property range. Accordingly, we investigated *DrugDiff*’s ability to respond to guidance by a property predictor (for details, see methods). For each property, we first generated 10,000 molecules *without* guidance to use their property distribution as a point of reference. We then introduced guidance to the generation process for both directions, i.e. to increase and to decrease the property. To further illustrate the responsiveness of *DrugDiff* to the guidance signal, we used seven different guidance strengths when guiding towards an increased or decreased property value, generating 10,000 molecules in each case. Guidance strength refers to the factor by which the predictor loss is amplified and guidance strength of zero represents generation without guidance. Figure 3 illustrates the property distributions of the generated molecules at different guidance strengths and for different molecular properties. The properties that were investigated here are (from left to right and top to bottom): logP, synthetic accessibility, quantitative estimation of drug-likeness, penalised logP, molecular weight, molar refractivity, number of cycles (i.e. rings), number of hydrogen bond donors, formal charge, number of rotatable bonds, number of hydrogen bond acceptors and the topological polar surface area. For continuous-valued properties, clear gradual shifts in density can be observed for increasing guidance strengths in both directions. Discrete properties are illustrated as stacked bar plots, with lower and higher property values clearly enriched for strong guidance to decrease and increase the property. A random molecule was selected and displayed for each set of molecules generated with the strongest guidance strength in both directions, additionally visualising the observed trends on concrete examples. For instance, when increasing the logP, i.e. guiding the generation towards more lipophilic molecules, the randomly selected molecule exclusively comprises carbon and hydrogen, whereas the molecule picked from the distribution with guidance toward low logP also includes several oxygen and nitrogen atoms as well as charges. When guiding for high and low numbers of rings, the randomly selected molecules contain six and zero rings, respectively. In addition to the evident response to the provided property guidance, also some anticipated confounders between properties become clear. For instance, increasing the number of rotatable bonds expectedly results in a general increase in the molecules’ size. For multi-conditional setups such implicit control over other properties should be considered to optimise the predictor panel in terms of fewer predictors to cover the desired set of properties.

**Figure 3:**
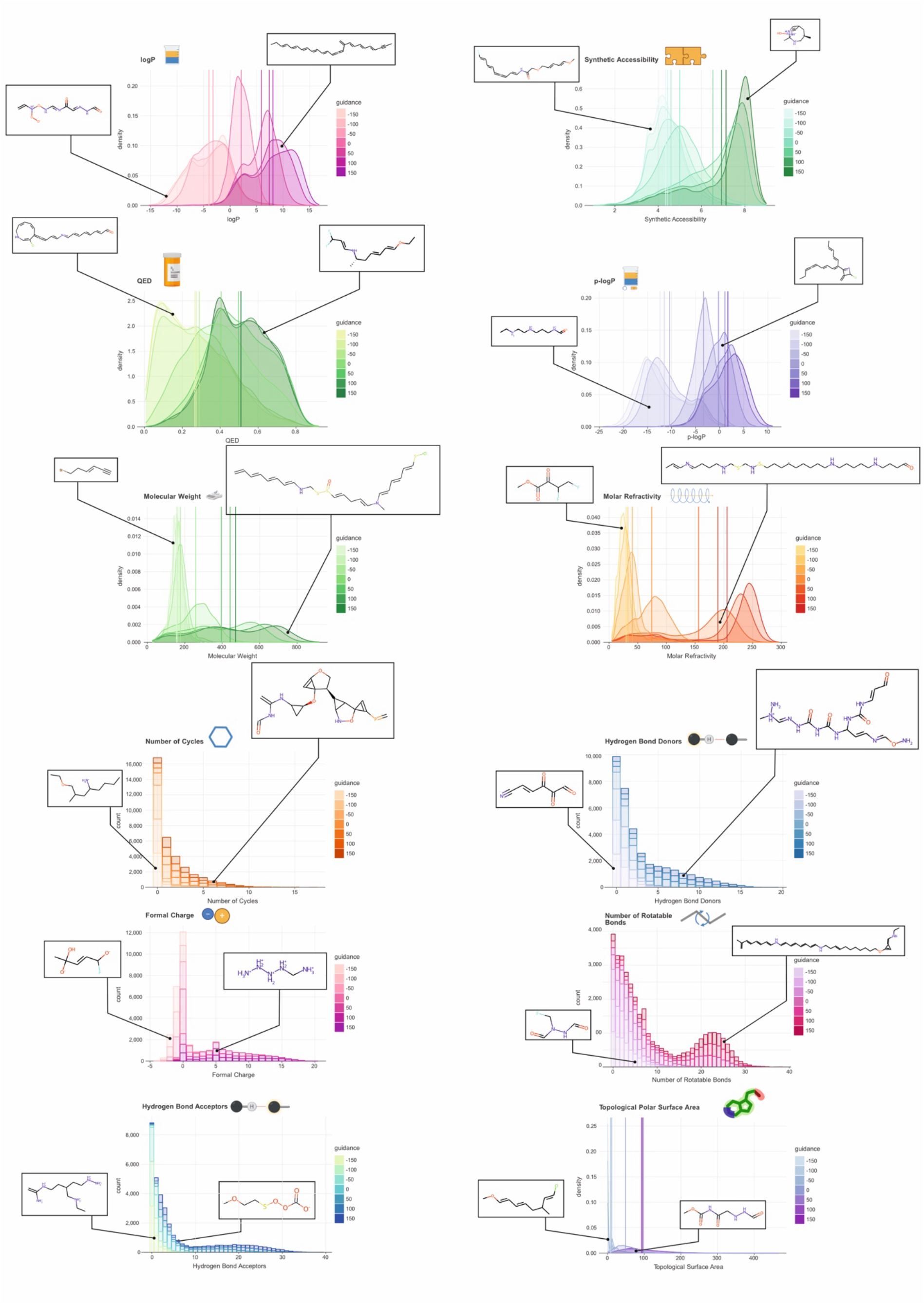
Single-Property Guidance. Shown are the distributions of various molecular properties in a set of 10,000 generated molecules when guiding the generation process with different guidance strengths. A guidance strength of zero equates to unguided generation, positive guidance strengths intend to shift the distribution towards higher values and negative guidance strengths towards lower values. Vertical lines represent the mean property value for each guidance strength. For illustration purposes, a generated molecule has been selected at random for strong positive and strong negative guidance and plotted alongside the distribution.

### Multi-Property Guidance

During drug development, it is often of particular interest to not only modify a single molecular property, but instead manipulate multiple properties in parallel. This task may include increasing or decreasing the different properties together or steering their values into opposing directions, reducing one while increasing another. We therefore next investigated *DrugDiff*’s ability to accommodate signals of several property predictors during the generation process for the purpose of multi-property guidance. Specifically, we considered the logP as well as the number of heavy atoms and therefore the overall size of the molecules. We considered four scenarios: 1) low logP and low number of atoms, 2) high logP and low number of atoms, 3) low logP and high number of atoms and 4) high logP and high number of atoms. For each scenario we generated 10,000 molecules. As can be seen in Figure 4A, the density of molecules generated in scenario 1 (teal) is indeed highest in regions of low logP and low number of atoms. Similarly, molecules generated in scenario 2 (orange) show much higher, positive logP values than those in scenario 1, while their number of atoms remains low. While in scenario 4 (green), a small subset is mislocated at low number of atoms the majority of molecules indeed have both high logP and number of atoms. Finally, scenario 3 (blue) — low logP and high number of atoms — was the hardest for the model to realise. While there is a population that fulfils these conditions, the large majority is located at lower heavy atom counts. Figure 4B shows a principal component analysis (PCA) with the first two principal components (PCs) displayed for the molecules generated in the different scenarios. The PCA was computed on the molecules’ MACCS fingerprints (rdkit v2022.09.5), which indicate the presence of predefined substructures in a molecule. Clear spatial separation can be observed between the molecules generated for the different scenarios, with PC1 separating lipophilic from hydrophilic molecules and PC2 separating small from large.

**Figure 4:**
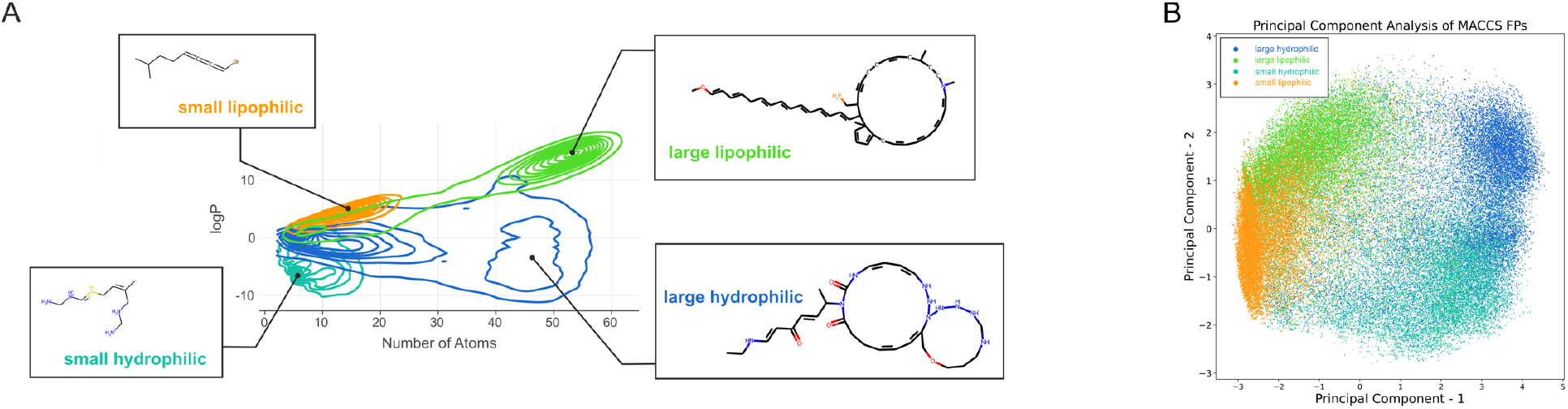
Multi-Property Guidance. A) Shown are the distributions of molecules in terms of logP and the number of heavy atoms generated with dual guidance in four different scenarios: high logP and low number of atoms (orange), low logP and low number of atoms (teal), high logP and high number of atoms (green), low logP and high number of atoms (blue). For each distribution, a molecule was picked at random and illustrated alongside it; B) Shown are the first two principal components of a PCA conducted on MACCS fingerprints of the molecules from the four aforementioned scenarios. The colour-coding corresponds to that in A).

## Discussion

In this work, we have introduced a latent diffusion model with property guidance for the generation of novel small molecules with desired target properties. This was motivated by the underrepresentation of small molecules in the recent surge of deep generative models — specifically diffusion models — in the molecular context, which has mostly focused on proteins, despite the prominent role of small molecules in the drug landscape. Furthermore, existing generative models for small molecules are rigid in their design and do not allow easy adaptation to different use cases. This particularly concerns the way molecular properties are included in the model design. While the majority of works include them as conditions during the training process, we decided against this approach since it requires retraining of the entire model whenever properties are to be exchanged or added. Instead, *DrugDiff* utilises predictor guidance, where only chemical property predictors need to be trained and plugged in to include additional properties but the generative model does not require retraining. Further, by using a latent diffusion model, the VAE can be easily replaced in the framework for one that handles a different modality than SELFIES, while the overall setup of the diffusion model and its training procedure remain unchanged. The conducted experiments clearly demonstrate that *DrugDiff* i) was able to learn the breadth of its training distribution, ii) can distinctly manipulate the molecular properties of molecules during the generation process, and iii) is capable to also expand this to a multi-property setting. In the context of multi-property guidance, it is important to highlight that the simultaneously applied properties must be reasonable. As could be seen on the example of decreasing logP — i.e. increasing hydrophilicity — and at the same time increasing the number of heavy atoms — i.e. making the molecules larger —, it was difficult to produce a large population of molecules that fulfilled both criteria. Given that large organic substances are predominantly made up of a lipophilic hydro-carbon backbone, difficulties in creating large organic molecules with very low logP are to be expected. Hence, when using multi-property guidance, the desired conditions must be chosen to fit realistic scenarios in order to ensure a high yield.

In summary, *DrugDiff* is a diffusion model for the generation of small molecules with desired target properties, which was build in a highly flexible framework. The modular architecture allows high customisability, allowing the exchange of the VAE including the molecular representations it receives, as well as the properties used for guidance, making it easily adaptable to diverse application scenarios while minimising the required retraining of the model. Future expansions should include more complex properties, such as target binding affinity, binding selectivity or systemic biological effects. Another future direction is training on much larger datasets to cover a vaster chemical space. While good results could already be achieved by training on the relatively small training set ZINC250K, more training data can be expected to further push the performance. Since the diffusion model learns from the VAE’s latent space, latent space quality is instrumental for the diffusion model’s success. Optimising the VAE with respect to the representation of chemistry in the latent space can therefore be expected to further improve performance. Additionally, a common task in drug development pipelines is substructure optimisation. Here, a substructure is provided as a starting point and modified to improve selected molecular properties. Incorporating this into the *DrugDiff* framework will add another layer of utility. Lastly, an end-to-end experiment starting with *in silico* generation of leads with *DrugDiff*, followed by synthesis and *in vitro* and/or *in vivo* evaluation will give insight into the exact speed-up of the development process offered by the model.

## Methods

### The Model

The model constitutes three parts, a variational autoencoder that maps SELFIES onto a latent representation, the diffusion model that is trained to generate such molecule latents and a collection of property-predictors that are used to guide the diffusion process.

### VAE

Variational Autoencoders [26] comprise two models, an encoder and a decoder. The encoder *g*(*x*) = *z* is used to embed higher-dimensional data onto a continuous-valued latent space of typically smaller size. The decoder *f*(*z*) = *x*′ ≈ *x* is trained to revert the embedding back to the original input. The VAE used for the embedding of molecules for further processing by the diffusion model was taken from Eckmann *et al*. [16]. The model was pre-trained on one-hot encodings of SELFIES from the ZINC250K dataset, a subset of the ZINC database [17], [18], [27], comprising approximately 250,000 small molecules that fulfil the Lipinski Rule of 5 [28]. The model utilises a latent size of 1024. Training the VAE was done following the steps described in the repository accompanying the paper [16].

### Diffusion Model

Our diffusion model is a *latent diffusion model* based on the implementation by Rombach et. al [29]. More precisely, the underlying model architecture is a *denoising diffusion probabilistic model* (*DDPM*) [30] with a linear layer, three fully connected residual blocks, a layer norm and another linear layer. Timesteps are embedded and provided to the fully connected residual blocks via feature-wise linear modulation layers.

*DDPMs* are trained to revert a Markov chain of noising steps, the diffusion process. This diffusion process incrementally adds Gaussian noise to a sample over 0, . . ., *T* timesteps such that a sample *x*_0_ from the original domain is transitioned into *x*_*T*_, which is Gaussian noise for large *T*. The noising step at *t* is conditioned on the noised sample *x*_*t*−1_ of the previous timestep

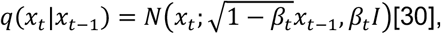

where *t* is taken from a pre-defined, time-dependent noising schedule. In this formulation, the computation of *x*_*t*_ first requires the computation of all previous timesteps *x*_0:*t*−1_, however, applying the reparameterization trick [26], [30], it can be directly conditioned on *x*_0_ with

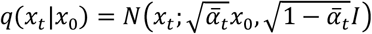

where *α*_*t*_ = 1 – *β*_*t*_ and 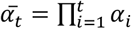. The noising schedule used here is linear. The *DDPM* is trained to reverse this diffusion process and therefore generate a sample from *q*(*x*_*t*−1_|*x*_*t*_). Stepwise application for *t* = *T*, . . ., 0 then allows the generation of samples from the original domain, *x*_0_ ∼ *q*(*x*_0_), starting from Gaussian noise *x*_*T*_. In this work, we are rephrasing the sampling step as a noise predictor that predicts *t* rather than *x*_*t*−1_ directly, as proposed by Ho *et al*. [30], which is trained using

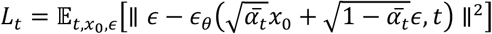

for 0 < *t* ≤ *T*. Note that 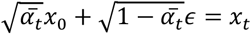 after the reparameterization trick. The latent diffusion model was trained for 100 epochs on the VAE-embeddings of the ZINC250K data.

### Property Predictors

We use a series of property predictors to guide the diffusion model during sampling. The property predictors followed the implementation of Eckmann *et al*. [16] and were trained on the ZINC250K dataset. Drug-likeness (QED), logP, molecular weight, molar refractivity, topological polar surface area, number of H-bond acceptors/-donors, number of rotatable bonds, number of atoms, formal charge and number of rings were all computed using *rdkit* (v2022.09.5). The synthetic accessibility was calculated using the *sascorer*.*py* from [16] which is based on[21]. The p-logP was computed as follows:

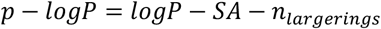

where *n*_*largerings*_ is the number of rings with more than 6 atoms. All predictors were trained on decoded one-hot encodings of the molecules. During guidance, the noisy latents were first decoded by the VAE’s decoder and then passed to the property predictors. We then computed the gradients based on the predicted properties and used them to manipulate the predicted noise *∈*_*t*_.

### Evaluation Metrics

To evaluate the molecules generated by our latent diffusion model, we used three metrics: 1) novelty, the fraction of generated molecules that are not found in the training set, to ensure that the model does not simply learn to copy the molecules; 2) uniqueness, the percentage of unique molecules among all generated ones; 3) internal diversity, a metric implemented in the *molsets* library that assesses how chemically diverse the generated molecules are. With this metric we can detect mode collapse, i.e. the model producing only a set of highly similar molecules to satisfy a given property.

## Ethics Note

An issue that needs addressing is the dual-use problem of models like the one presented here. Generating molecules that manifest desired properties has many beneficial use cases, for example in the medical context or materials sciences. However, it also bears the inherent risk of being misused for malicious intent, for example to generate molecules that maximise toxicity. The model described in this work is solely intended to be used in a beneficial context and in accordance with the law.

## Acknowledgments

This work was supported by the HGF Helmholtz AI grant Pro-Gene-Gen ZT-I-PF5-23 as well as the Helmholtz Association’s Initiative and Networking Fund on the HAICORE@FZJ partition. Further support came from the EU supported BMBF grant PriSyn (16KISA030). Figures were partially created using Biorender.com.

## Supplementary Information

**Supplementary Table 1:** GuacaMol distribution learning benchmark. VAE was re-trained on the GuacaMol benchmarking set, *DrugDiff* was then re-trained on the VAE latent space. The similarity of the benchmark results emphasise that *DrugDiff* is capable of fully learning the VAE’s latent space. KL: Kullback-Leibler Divergence. FCD: Fréchet ChemNet Distance.

|  | Validity | Uniqueness | Novelty | KL div | FCD |
| --- | --- | --- | --- | --- | --- |
| VAE | 1.0000 | 0.9851 | 0.9972 | 0.6578 | 0.0038 |
| DrugDiff | 1.0000 | 0.9884 | 0.9973 | 0.6609 | 0.0035 |

